# Scalable Bayesian inference for high-dimensional neural receptive fields

**DOI:** 10.1101/212217

**Authors:** Mikio C. Aoi, Jonathan W. Pillow

**Affiliations:** Princeton Neuroscience Institute Princeton University, Princeton, NJ 08544

## Abstract

We examine the problem of rapidly and efficiently estimating a neuron’s linear receptive field (RF) from responses to high-dimensional stimuli. This problem poses important statistical and computational challenges. Statistical challenges arise from the need for strong regularization when using correlated stimuli in high-dimensional parameter spaces, while computational challenges arise from extensive time and memory costs associated with evidence-optimization and inference in high-dimensional settings. Here we focus on novel methods for scaling up *automatic smoothness determination* (ASD), an empirical Bayesian method for RF estimation, to high-dimensional settings. First, we show that using a zero-padded Fourier domain representation and a “coarse-to-fine” evidence optimization strategy gives substantial improvements in speed and memory, while maintaining exact numerical accuracy. We then introduce a suite of scalable approximate methods that exploit Kronecker and Toeplitz structure in the stimulus autocovariance, which can be related to the method of expected log-likelihoods [1]. When applied together, these methods reduce the cost of estimating an RF with tensor order *D* and *d* coefficients per tensor dimension from *O*(*d*^3*D*^) time and *O*(*d*^2*D*^) space for standard ASD to *O*(*Dd* log *d*) time and *O*(*Dd*) space. We show that evidence optimization for a linear RF with 160*K* coefficients using 5*K* samples of data can be carried out on a laptop in < 2s.

## 1 Introduction

For over a century scientists have characterized a neuron’s basic stimulus selectivity as a first step in analyzing its other properties [2]. This is usually achieved by solving a regression problem whereby a neuron’s linear receptive field (RF) is modeled as a noisy mapping from the space of high-dimensional stimuli to a scalar variable describing the neuron’s response (ex. spike counts, calcium fluorescence, membrane voltage, etc.). However, least-squares and maximum likelihood estimates of the RF exhibit poor statistical performance in high-dimensional settings, requiring strong regularization [3]. This regularization is especially necessary when stimuli display correlated structure.

Bayesian methods offer a solution to the statistical problem by using prior information to regularize RF estimates. Ridge regression, the simplest such method, merely biases the estimated coefficients towards zero. Recent advances have focused on priors that exploit known features of RF structure, such as smoothness and sparsity, which give substantial improvements in statistical efficiency [4, 5, 6], potentially reducing the time and costs of conducting laboratory experiments. However, these methods become intractable as stimulus dimensionality grows large. In particular, for a d-dimensional stimulus the empirical Bayes estimator requires 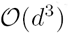 time and 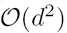 memory per evaluation of the marginal likelihood, making the Bayesian framework prohibitive for large *d*. Since this computational bottleneck can be localized to operations on covariance matrices, we focus our efforts on identifying computational savings for operations in both the *prior* covariance and the *stimulus* covariance.

In this paper, we describe strategies for making empirical Bayes RF estimation tractable in high dimensions and large sample sizes. We focus on the *automatic smoothness determination* (ASD) estimator [5], though our results have applicability to a variety of models based on conditionally Gaussian priors. We demonstrate these methods using both simulated and real neuronal data with high-dimensional stimuli.

## 2 The ASD regression model

In this section we outline the basic elements of the ASD model of Sahani and Linden [5] and point out some of the factors making it resistant to scalabilty.

The ASD model assumes that a neuron's response at time *t*, *y*_*t*_, to the stimulus 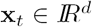 is a linear function of the stimulus corrupted by iid noise:

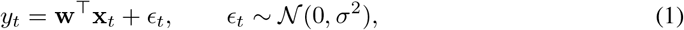

where 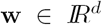 is the RF and 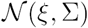 represents the Gaussian distribution with mean ξ and covariance Σ. The log-likelihood is therefore given by

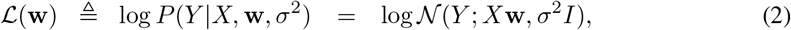

where *X* = (x_1_, x_2_,…, x_*N*_)^*T*^ is the design matrix with stimuli for each trial along its rows, and *Y =* (*y*_1_,…, *y*_N_)^⊤^ is the vector of responses to each stimulus.

The key to the ASD model's efficiency is the use of a Gaussian process (GP) prior over the weights, 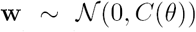, with covariance matrix *C*(*θ*) defined by [*C*(*θ*)]_*jk*_ = *ρ* exp(-\z_*j*_ — z_*k*_|^2^/2*ℓ*^2^), where hyperparameters *θ* = {*ℓ*, *ρ*} include a length scale *ℓ* controlling smoothness and a marginal variance *ρ* controlling magnitude, and *Z*_*j*_, *z*_*k*_ denote the location of RF elements in spatiotemporal pixel space. This covariance function is commonly referred to as the squared exponential^1^ (SE) in the machine learning literature [8].

The ASD model weights can be learned by an empirical Bayes procedure [9] (also known as type-2 maximum likelihood [10] and the evidence approximation [11, 12]) which involves 2 steps. First, the hyperparameters *θ* and noise variance *σ*^2^ are set by maximizing the marginal likelihood (or “evidence”) [13], which is the probability of the data given the hyperparameters,

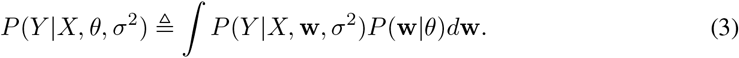

For the Gaussian likelihood and prior of the ASD model the log evidence has a convenient closed-form expression;

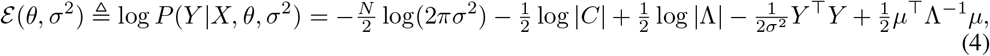

where 

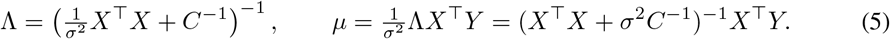

are the posterior covariance and mean, respectively. Second, the RF estimate is given by 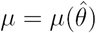, the MAP estimate of w conditioned on the optimal values of *θ* and *σ*^2^.

This estimator has excellent performance in many settings and improves estimation accuracy and generalizability of RF estimates compared to maximum likelihood [5]. Empirical Bayes is often adopted as a computationally efficient alternative to MCMC or cross validation methods.

### 2.1 Obstacles to scalability

While the ASD model is well known, it has not been used in high-dimensional settings due to the poor scaling of the method when computed by brute force. The computational bottleneck comes about from the fact that each evaluation of the log evidence (4) requires the calculation of both the inverse and determinant of Λ, each of which requires 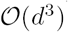 operations, where *d* is the number of parameters. In the following sections we present several approaches to efficient evaluation of the evidence function, some exact and some approximate, and we will examine the impact that these methods have both on MAP inference and evidence optimization. We emphasize that these are *not* methods for efficient GP regression but are methods for efficient inference of a GP-regularized linear regression model.

## 3 Fourier-domain ASD

In this section we briefly outline the Fourier representation of a GP and show that this representation allows for a significantly reduced dimension of the ASD model without appreciable loss in accuracy. A more detailed discussion of the spectral representation of GPs is discussed elsewhere [14, 15, 16].

### 3.1 Padded Fourier representation of the ASD prior covariance

The prior covariance matrix *C* of the ASD model can be diagonalized by the Fourier transform, but not without some modification. Here we describe that modification and its basic intuition.

The ASD prior covariance can be represented in frequency coordinates by 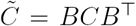 and the frequency-domain stimulus can be expressed as 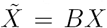, where *B* is the orthonormal discrete Fourier transform^2^ (DFT) matrix with 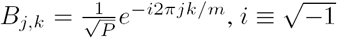.

The ASD model uses a *stationary* GP for a prior on the regression weights. Covariance kernels of stationary GP covariances have real, symmetric Fourier transforms and translate into making the matrix 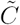 asymptotically diagonal where, if the Fourier transform of the GP kernel is given by 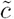, then the matrix 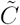 can be obtained by placing 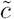 on the main diagonal with all other entries set to zero. This property allows for significant computational savings [6, 17, 18, 15, 14]. However, the covariance matrix of a finite observation of a stationary process is not directly diagonalizable. This is because, for the finite DFT, the diagonal Fourier representation corresponds to a stochastic process on a *periodic* domain.

In the context of the ASD model a periodic domain would imply that a RF had strong correlations across left and right (or top/bottom) edges, which is unlikely to be the case in practice. In order to take advantage of a diagonal Fourier domain covariance while avoiding spurious correlations over a circular boundary we can define our problem on a larger domain than that which is actually observed but with periodic boundaries. The basic idea is illustrated in Fig. **1A**.

Figure 1A (left panel) shows a typical prior covariance for a 1D domain with *d =* 200. As it is, this covariance matrix is not diagonal in Fourier coordinates. However, we can view this covariance as a sub-matrix of a circulant matrix defined on a somewhat larger, but periodic, domain (Fig. 1A, middle panel, note the spurious correlations at the top right and bottom left corners). This “extended” matrix is diagonal in Fourier coordinates. (Fig. 1B, right panel). In practice, one should choose to extend the spatial domain sufficiently to avoid appreciable correlations in the ASD prior sub matrix from the periodic boundary (i.e. the shaded area in Fig. 1A should be large enough to avoid the boundary effects to influence the true covariance). Put simply, we effectively pad the RF with enough zeros to avoid correlations between the edges of the RF. How large the barrier should be will depend on the the length scale of correlations in the GP kernel.

In our case, for the squared exponential kernel with length scale l, a sufficient boundary extension was achieved by adding 3l extra coefficients, giving a extended frequency domain representation of size 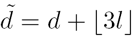, where 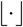 is the floor function. Note that this padding is virtual, in that it is never implemented in practice. Instead is it used to define a DFT matrix of appropriate dimensions for transforming the spatial-domain stimulus *X.* Because the size of the spatial representation determines the size of the frequency representation, the extended domain determines which frequencies should be represented by 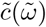. Indeed, the appropriate DFT matrix need not ever be formed, since *X* can be transformed by the 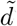-point FFT. The resulting representation can be used to conduct all inference procedures in precisely the same way as the spatial-domain problem.

**Figure 1:**
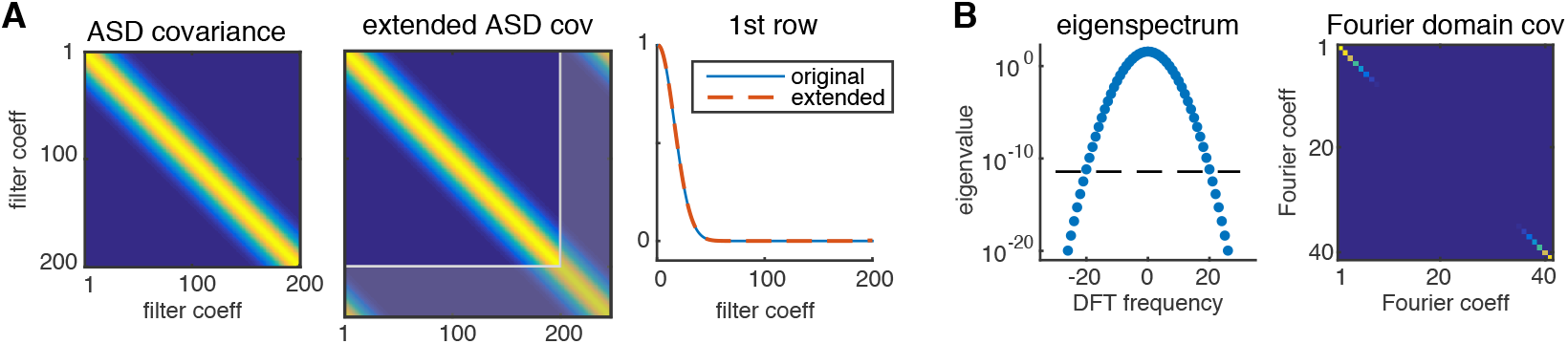
Efficient representation of ASD covariance. **(A)** Standard ASD covariance matrix for 1D (vector) filter with 200 coefficients, with length scale *l* = 15, and circularly extended (circulant) version with (200+3*l*)=245 coefficients. (Gray region corresponds to “unobserved” filter components). **(B)** The circularly extended covaraince has analytically computable eigenspectrum (left), allowing for efficient pruning of unnecessary dimensions, and is diagonalized by the Fourier transform (right).

### 3.2 Truncated spectrum

In order to ensure a diagonal Fourier domain prior covariance we extended the effective spatial extent of the filter, making the dimensionality of the problem larger than when we started. However, the ASD kernel has “low pass” character, indicating that there are many more frequencies represented than we actually need. The SE kernel for example has the Fourier representation 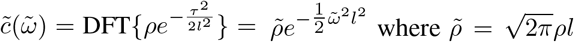 is the frequency-domain variance and 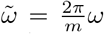 are the “effective frequencies”. However, note that c decays quickly as the frequencies become large, with faster decay occurring as *l* increases. Figure 1B (left) shows the decay of the Fourier coefficients on the log scale for *l* = 15 with a filter length of 200. Thus, the Fourier-domain filter coefficients at high frequencies will tend to be small, suggesting that they may be safely disregarded. For a principled rule for choosing which coefficients to ignore, one could set a threshold to maintain the condition number of 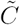 below some value *δ*. Again, because we have an analytical expression for 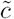 we can explicitly define this threshold for a given length scale for the SE kernel to be

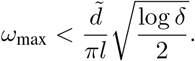

Cleary, the truncated representation is more efficient, and more accurate, as the length scale *l* increases. Importantly, when the condition threshold is set high enough spectral ASD can be exact up to machine precision, even though the dimension is reduced substantially.

The dimension-reducing effect of this thresholding can be dramatic, as is illustrated in the example in Fig. 1B. For the threshold of *δ* = 10^8^ (chosen to stay well — within floating-point accuracy), we achieved a ≈ 5 × reduction in the size of the representation (reducing the computational complexity by 125 ×).

The computational savings are realized by reducing the number of frequencies for 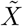 as well. Noting that most of the high-cost computations are performed on the matrix 

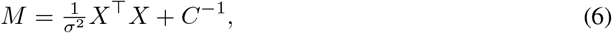

we find that the truncated Fourier setting reduces *M* from by a degree governed by *δ.* This dimension reduction principle applies for a 2D domain as well where, rather than setting the maximum frequency, we set the spectral radius.

#### 3.2.1 Course-to-fine parameter search

Because the Fourier representation of the ASD model becomes more computationally efficient as the length scale *l* of the prior covariance becomes large, it would be prudent to prioritize the optimization algorithm to search over larger length scales first. This principle suggests a “course-to-fine” strategy where we set the lower-bound on *l* fairly large to take advantage of computational efficiency in early iterations of the optimization. When the optimization bumps up agains the lower bound, that bound can be lowered and can be progressively decreased as needed, thereby ensuring that computation is not wasted on searching very small values of *l* if it is not necessary.

## 4 Structured approximate estimators for extreme scalability

Even in the case where we can reduce the dimensionality of the ASD model by Fourier approximation, the number of parameters may still be too large to make practical calculations. In particular, although we are able to diagonalize the prior covariance *C,* the sample covariance *X* ^⊤^ *X* is still *d′* × *d′,* where *d′* is the size of the reduced Fourier representation. This will mean that calculation of A and its log determinant will still be prohibitive as they scale as 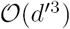.

One way for us to further reduce the computational complexity of this problem is to replace *X* ^⊤^ *X* with a structured estimator of its expectation. The intuition for this approach follows from the same logic as the expected log-likelihood (ELL) approximation [19, 1].

The argument is as follows: If the stimulus covariance is given by 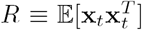 (assuming wlog 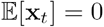) then as the number of samples *N* becomes large we have 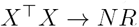. Thus, in settings with large *N* a law of large numbers argument can be made to approximate the likelihood with *N R* in place of *X* ^⊤^ *X* [19, 1]. The resulting cost function is called the *expected log-likelihood* and is given by

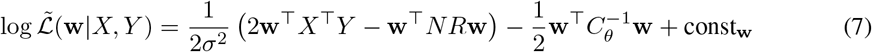

which is simply the likelihood in (2) with *X* ^⊤^ *X* replaced by *NR.* Maximizing the ELL with respect to w yields the maximum ELL estimator (MELE). If *R* has special structure then this can be exploited for computational efficiency.

Note however that we cannot simply use the ELL trick “out of the box” when we do not know *R* a priori. This is especially true when we are interested in studying neuronal responses to naturalistic stimuli, for which neither the spatio-temporal covariance, nor the relevant spatio-temporal resolution are well established. In this case, we can employ similar reasoning behind the ELL estimator to justify replacement of *X* ^⊤^ *X* with a structured estimator of *R.* The advantage of this approach is that we need not have detailed knowledge of the statistics of the stimulus, only that we know that its covariance has a certain properties.

Two properties that we will consider here are Toeplitz structure and Kronecker factorization. We next outline each of these properties and show how they provide computational savings.

### 4.1 Kronecker structure in *R*

Covariance matrices of tensor-structured data admit a Kronecker factorization which makes for convenient computational savings [20, 7]. An example of a tensor-structured RF would be a spatial filter which has two space dimensions (i.e. a 2D tensor).

For tensor-structured RF weights and stimulus, the prior covariance *C* and the stimulus covariance *R* can be represented as the Kronecker products of smaller matrices. For example, for tensor dimension 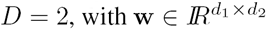, the prior covariance of w has dimensions (*d*_1_*d*_2_) × (*d*_1_*d*_2_) and given by 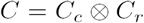, where *C*_*c*_ and *C*_*r*_ are the *d*_2_ x *d*_2_ column covariance, and *d*_1_ x *d*_1_ row covariance, respectively.

Empirical Bayes estimation for the ASD model requires calculations involving both the evidence and posterior with operations on the matrix 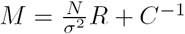. In this case,

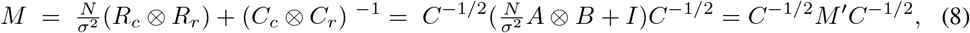

where 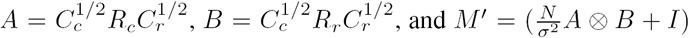 are real, symmetric and positive definite. Therefore, *A* and *B* have eigen-decompositions *A* = *UPU*^*⊤*^ and *B = VQV*^⊤^, where *U, V* are orthonormal and P, *Q* are diagonal, respectively. This allows us to obtain the eigen-decomposition for *M':*

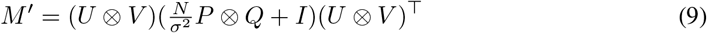

where (*U* ⊗ *V*) is orthonormal and 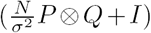 is diagonal [21]. This greatly simplifies operations on *M.* In particular, the inverse and determinant of *M'*:

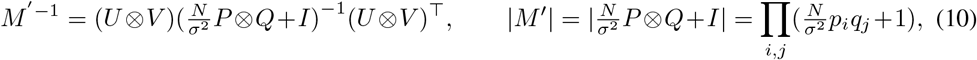

where *p*_*i*_ and *q*_*j*_ denote the diagonal entries of *Q* and *P.*

Thus, the the *D =* 2 case the Kronecker property has the effect that inversion and matrix determinant calculations are now bought at the cost of calculating the eigenvalues/vectors of *A* and *B.* In fact, the *d*_1_*d*_2_ x *d*_1_*d*_2_ Kronecker products never have to be formed and all operations can be performed on using only the Kronecker factors, allowing for substantial memory savings as well. These tricks become more efficient as the tensor dimension grows. In general, for tensor dimension *D,* and array dimension *d*_*p*_ along tensor dimension *p,* the Kronecker factorization allows us to reduce the computational and memory costs from 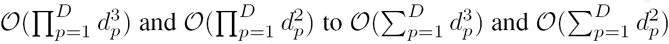, respectively. Furthermore, matrix-vector multiplications, where matrices have Kronecker structure, can be implemented by efficient algorithms [20], allowing for further computational savings.

### 4.2 Toeplitz structure

If the stimulus distribution is stationary, then its covariance will be Toeplitz. A Toeplitz matrix *R* may be represented as a 1D autocovariance sequence *r*(*τ*), rather than a 2D matrix. Each row of *R* is therefore a shifted copy of *r*(*τ*). The ASD prior covariance (Fig. 1, left) displays exactly this kind of structure. If *R* is indeed Toeplitz, then it is subject to the same diagonal Fourier representation as *C* (with the same caveat that diagonalization requires an extended representation as described in Section 3.1). Therefore, since the same basis will diagonalize both *R* and *C*, the *M* matrix defined in (8) will have the representation

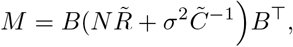

where both 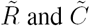 are diagonal. Again, we need not ever apply the matrix multiplication of *B* (the DFT matrix) or its inverse (*B*^*T*^) since we can simply apply the FFT and its inverse to the first row of *R* to obtain the Fourier coefficients.

Furthermore, provided that *R* has a “low-pass” character, we can apply all of the same tricks for implementing a Fourier-domain dimensionality reduction to *R* as we did in Section 3.1 for *C,* reducing the complexity of all operations on *M* accordingly. Thus, using the Toeplitz approximation reduces the complexity of estimating an RF with tensor dimension *D* from 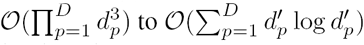, where 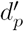 is the reduced array dimension corresponding to thresholding the spectral norm.

### 4.3 ELL with approximate R

An important feature of both the Kronecker and Toeplitz strategies of the ELL approximation is that both of these properties can be exploited even when the true stimulus covariance *R* is unknown, such as when the stimulus is given by customized patterns or naturalistic scenes. For the Kronecker method, *R*_*c*_ and *R*_*r*_ can be estimated from the data and used as *plug-in* estimators of *R.* Similarly, the Toeplitz methods can be implemented with a sample autocovariance sequence *r*(*τ*).

## 5 Experiments

The empirical Bayes framework for the ASD model is implemented in two stages, evidence optimization (for hyperparameters *θ*) and MAP estimation (for filter parameters w). Both the evaluation of the evidence and filter estimation require operations with complexity 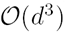 and the methods described above may be used to make both computations tractable. In the interest of assessing the impact of the approximate approaches on each stage, we will examine them separately and then provide examples of empirical Bayes estimation of a large-scale RF for both real and simulated data.

**Figure 2:**
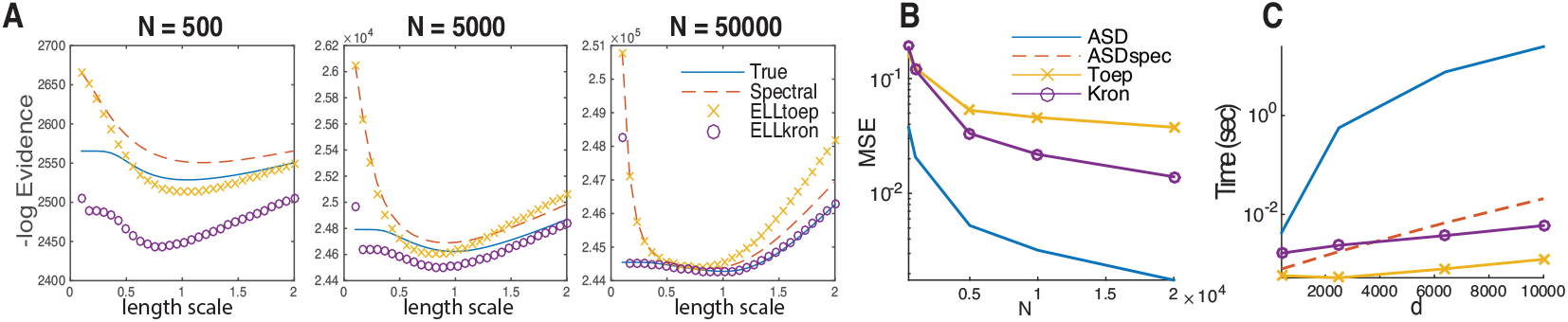
Quality of structured ELL approximations. **A**) Convergence of the log evidence for 2D filter as a function of the length scale *l*. A 20 x 20 filter was drawn with *σ*^2^ = 1000. The true length scale is *l* = 1. **B**) MSE of a 80 x 80 filter as a function of sample size and the time for calculation as a function of filter dimension for a sample of 5000 observations. C) Wallclock time for estimating filters of specified dimension *d.* Metrics in **B** and **C** are averaged over 30 trials.

### 5.1 Evidence Approximation

We evaluated log evidence for data generated for a 20 × 20 filter sampled from a GP with *l* = 1 and, noise variance *σ*^2^ = 1000. Figure 2A shows the negative log-evidence for the true, spectral, Toeplitz ELL and Kronecker ELL methods as a function of *l* over samples sizes ranging three orders of magnitude. In the empirical Bayes framework this function is minimized in order to identify optimal hyperparameters and therefore *lower* values of the negative log-evidence indicate *higher* evidence. The spectral and ELL methods appear to be biased in opposing directions with respect to the true evidence. Specifically, the spectral methods are biased upwards, while the ELL approximations are biased downwards. This may be because the ELL approximation assumes more information about the filter than is warranted due to the use of the estimated stimulus covariance in place of the the sample stimulus covariance. The spectral method on the other hand correctly biases the negative log-evidence upward to reflect the reduced information due to reduced stimulus dimensions. However, the spectral approximations appear to be more strongly convex than either the ELL or the true evidence, suggesting that the spectral methods may be better conditioned in practice for evidence optimization. This improved resolution may be due to the low-pass properties of the dimension reduction that may “filter out” information that is not important to the hyper parameters.

The three methods clearly appear to converge to the true evidence as the sample size increases, particularly in the neighborhood of the true length scale (*l* = 1). However, they all appear to achieve their minimum at a length scale that slightly deviates from the true *l*, depending on the sample size. While the true evidence is extremized at *l* = 1, the Kronecker approximation appears to be extremized at a slightly smaller value of *l* for smaller sample sizes (Fig. 2 left panel), while the spectral methods are relatively unbiased. However, for larger sample sizes, the Kronecker approximation converges to the true evidence for all *l*, while the spectral methods appear to be slightly biased. A deeper study of this behavior is needed in order to understand why this occurs and to determine the appropriateness of each method in a given sampling regime.

### 5.2 MAP estimation by empirical Bayes

To compare each method for filter estimation we generated 5K corrupted responses *y* simulated with a 80 x 80 Gabor filter with noise variance *σ*^2^ = 125. We generated random stimulus vectors x from a GP with *l* = 1.5 and *ρ =* 2. Figure 3A shows results for filter estimation for the three methods presented, demonstrating that they have comparable quality. Figure 2B shows the mean-squared error (MSE) of the estimated 80 x 80 filter as a function of the number of samples and Fig. 2C shows the wallclock time for calculation as a function of the number of filter coefficients for *N =* 5000 samples. Note that the spectral representation for ASD has nearly equal MSE with the conventional estimator but is orders of magnitude faster and scales more favorably. The approximate methods however (Toeplitz and Kronecker) admit large gains in speed over the unstructured MAP estimator, and scale more modestly than the either the space-domain or spectral domain MAP estimates, but at the cost of increased MSE. It is notable however, that the MSE of even the Toeplitz approximation, which has the largest MSE of the methods presented, is only ≈ 9 % the size of the filter variance. We notice that for small *d* the spectral MAP method is actually faster than the Toeplitz method. This may be due to the additional cost of bookkeeping incurred by the diagonalization of *R.* However, since this cost is small, the Toeplitz estimator maintains extremely low computational cost even at very high dimensions.

**Figure 3:**
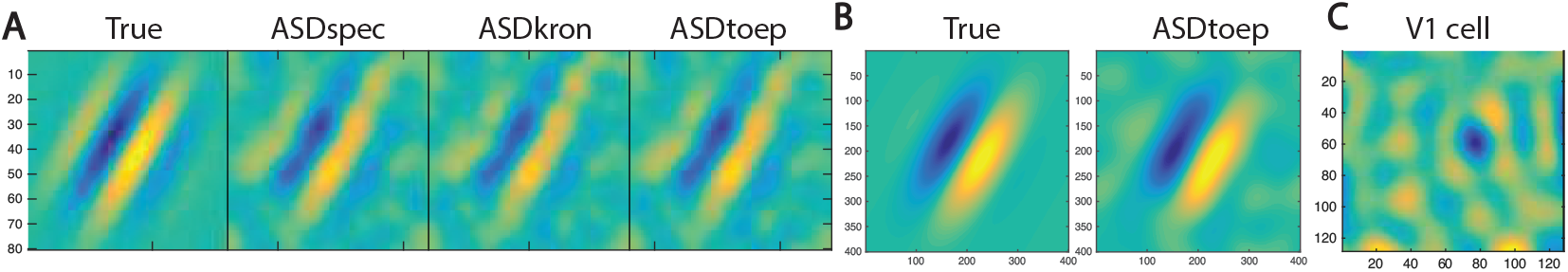
MAP estimation of 2D Gabor filter. **A**) Estimation of 80×80 filter using the specified methods. **B**) Estimation of 400×400 filter using the Toeplitz method. **C**) Estimated receptive field from the calcium signal of a V1 cell using a 128×128 stimulus by the spectral method.

We note here that all of the times that we assess are clocked *after* the calculation of sufficient statistics, where calculation of the stimulus covariance may require considerable time for very large sample sizes and dimension. However, such a calculation only has to be performed once and all inferential procedures can be iterated using the same set of sufficient statistics. Therefore, for our analyses we assess the time required to evaluate the filter estimator or the evidence.

#### 5.2.1 Large-scale empirical Bayes

We generated 5 thousand samples of a response from a 400×400 Gabor filter giving a total filter size of 160K coefficients. Note that there are no reference hyperparameters for this filter since it is not a realization of a GP and therefore is a model mis-fit. We instead identified optimal hyperparameters by maximizing the evidence and estimated the filter using the Toeplitz ELL estimator. Also note that the full stimulus covariance has 25.6 billion elements; more than can be represented on a desktop computer with double precision accuracy with current technology. The result is shown in Figure 3B.

The spectral representation of the filter and Toeplitz estimate of the stimulus covariance reduced the representation from 160K to 289 parameters. After calculation of sufficient statistics was completed, the empirical Bayes estimate of the filter was calculated in 1.97 seconds. The resulting estimate (Fig. 3B) had a error variance that was 4.12% the magnitude of the filter variance.

#### 5.2.2 Scalable ASD estimation of a neuronal RF

To demonstrate our spectral ASD method on real data we estimated the receptive field of a mouse V1 neuron (Fig. 3C). The stimulus was 128×128 and had 5600 stimulus presentations. Note that, due to the size of the stimulus (16,384 coefficients) whitening of this RF would have been impossible using conventional methods. Using the spectral method presented here, estimation of this RF, including calculation of sufficient statistics, was performed in ≈ 27 seconds on a laptop computer. Notably, we did not have to have detailed knowledge of the stimulus statistics, nor did we know the appropriate resolution with which to estimate the RF (to inform ad hoc spatial down-sampling).

## 6 Conclusions

We have outlined three basic strategies for scalable Bayesian RF estimation:

1. **Spectral representation:** allows for a diagonal representation of the prior covariance
2. **Kronecker plugin estimator:** allows for Kronecker representation of posterior covariance, which reduces all matrix calculations to the size of Kronecker factors.
3. **Toeplitz plugin estimator:** allows for diagonal representation of the posterior covariance.

All three strategies yielded considerable savings in computation and memory and all implementations could easily carry out filter estimation of over ten thousand coefficients within seconds on a laptop computer. We hope that the methods provided here will allow neuroscientists to consider large-scale RF estimation problems that may have been previously precluded due to computational costs. Further improvements in scalability may be achieved by combining the favorable properties of both the spectral methods and the Kronecker method when the conditions are appropriate. Identifying these conditions and how to implement them are the subject of current work in our lab.

## Footnotes

1 Although we deal exclusively with the squared exponential kernel in this paper, the methods are applicable to any stationary covariance with analytical Fourier transform including the Matérn kernel as well as more expressive mixture kernels like those found in [7].

2 The m-point discrete Fourier transform has frequencies 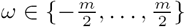.

